# The single mitochondrion of the kinetoplastid parasite *Crithidia fasciculata* is a dynamic network

**DOI:** 10.1101/388660

**Authors:** John DiMaio, Gordon Ruthel, Joshua J. Cannon, Madeline F. Malfara, Megan L. Povelones

## Abstract

Mitochondria are central organelles in cellular metabolism. Their structure is highly dynamic, allowing them to adapt to different energy requirements, to be partitioned during cell division, and to maintain functionality. Mitochondrial dynamics, including membrane fusion and fission reactions, are well studied in yeast and mammals but it is not known if these processes are conserved throughout eukaryotic evolution. Kinetoplastid parasites are some of the earliest-diverging eukaryotes to retain a mitochondrion. Each cell has only a single mitochondrial organelle, making them an interesting model for the role of dynamics in controlling mitochondrial architecture. We have investigated the mitochondrial division cycle in the kinetoplastid *Crithidia fasciculata*. The majority of mitochondrial biogenesis occurs during the G1 phase of the cell cycle, and the mitochondrion is divided symmetrically in a process coincident with cytokinesis. Mitochondrial division was not inhibited by the putative dynamin inhibitor mdivi-1, although mitochondrial membrane potential and cell size were affected. Live cell imaging revealed that the mitochondrion is highly dynamic, with frequent changes in the topology of the branched network. These remodeling reactions include tubule fission, fusion, and sliding, as well as new tubule formation. We hypothesize that the function of this dynamic remodeling is to homogenize mitochondrial contents and to facilitate rapid transport of mitochondria-encoded gene products from the area containing the mitochondrial nucleoid to other parts of the organelle.

## Introduction

Mitochondria are essential organelles for most eukaryotic cells. A typical cell contains a dynamic population of mitochondria whose architecture and distribution are maintained by protein-mediated mechanisms including fusion and fission events, mitochondrial movement, and positional tethering [1, 2]. The balance between membrane fusion and fission reactions, collectively called mitochondrial dynamics, is of particular importance as it contributes to several essential functions [3]. First, mitochondrial dynamics helps to establish mitochondrial morphology, which is closely linked to function [4–6]. Elaborate mitochondrial networks are associated with high rates of aerobic respiration, while more simplified mitochondria occur in cells with low respiratory activity [7]. Mitochondrial dynamics are also coordinated with the cell cycle in order to ensure the even distribution of mitochondria into daughter cells [8–10]. Fusion homogenizes proteins and other macromolecules which is important for overall mitochondrial health and network function [11–14]. In contrast, fission is increased during mitosis to allow the stochastic partitioning of organelles into daughter cells. Finally, mitochondrial dynamics allows for organelle quality control [15, 16] by rescuing [17–19] or disposing [3, 20] of mitochondrial fragments produced by depolarization of the membrane potential.

Mitochondrial fission and fusion are mediated by distinct members of the dynamin GTPase superfamily, and inhibition of these activities produces distinct morphological phenotypes. A lack of fission produces a more highly-fused, interconnected mitochondrial network, while interrupting fusion results in small, fragmented mitochondria [21–23]. Many other proteins have been implicated in mitochondrial dynamics [1, 6, 21, 24]. These studies have used model systems such as *Saccharomyces cerevisiae, Drosophila melanogaster*, and mammalian cells [25]. Much less is known about how and if these processes occur in other eukaryotic organisms. Kinetoplastid parasites are unusual among eukaryotes in that each cell contains a single mitochondrion. Thus, it is essential that mitochondrial biogenesis and division are coordinated with the cell cycle. This is also true for other organisms with single mitochondria, such as *Toxoplasma gondii* [26]. For example, in *Trypanosoma brucei*, mitochondrial fission mediates the final division event that splits one mitochondrion into two immediately prior to cytokinesis. This process is mediated by a dynamin-like protein called TbDLP. In other eukaryotes, dynamins that function in mitochondrial dynamics differ from the classical dynamins that mediate vesicle fission during endocytosis. Interestingly, *T. brucei* and other kinetoplastids lack classical dynamins [27, 28]. In fact, most kinetoplastids encode a single DLP, suggesting that a single enzyme can function in both mitochondrial fission and endocytosis, as has been demonstrated for bloodstream form *T. brucei* [29, 30]. Furthermore, kinetoplastid genomes lack identifiable orthologs for most other mitochondrial dynamics proteins, leading some to conclude that conventional fission and fusion outside of organelle division do not occur in these organisms [30, 31]. However, mitochondrial dynamics has been demonstrated in plants despite a lack of orthologs for proteins expected to mediate these processes [32].

We are interested in the inherent properties of mitochondrial networks and in exploring the unique challenges faced by eukaryotic organisms with a single mitochondrion and mitochondrial nucleoid. For this, we decided to work with the model kinetoplastid *Crithidia fasciculata*, which is a monoxenous parasite of mosquitoes. These parasites live on flowers, fruit, or in aquatic environments before being taken up by a mosquito [33]. *C. fasciculata* presents several practical advantages for investigating kinetoplastid cell biology. It can be grown in large quantities, it is genetically tractable, and its cell cycle can be easily synchronized. They have two developmental forms, a swimming nectomonad and a non-motile haptomonad, both of which can be generated in culture [34]. The haptomonad stage is particularly well-suited for live-cell imaging.

Here we describe the mitochondrial division cycle in *C. fasciculata*. We found that mitochondrial biogenesis is coordinated with the cell cycle and occurs mainly in G1, while mitochondrial division occurs shortly before cytokinesis. We observed through examination of live cells that the mitochondrion is dynamic and is constantly remodeled through both fission and fusion events. Furthermore, we have visualized the complex process of mitochondrial DNA division in these cells. Collectively, these observations provide a new perspective on the kinetoplastid mitochondrion and reveal intriguing avenues of investigation. The study of mitochondrial dynamics in early-diverging eukaryotes provides an important evolutionary perspective on both the function and mechanisms of these processes and may have broad relevance for other eukaryotic systems.

## Materials and methods

### Parasite growth

*C. fasciculata* strain CfC1 was grown in brain heart infusion (BHI) medium supplemented with 20 µg/ml bovine hemin (Sigma) at 27 °C. Cells grown on a rocker were passaged as nectomonads every 2-3 days to maintain a density between 10^5^ and 10^8^ cells/ml. Cell densities were determined by mixing a small sample of cells with an equal volume of 3% formalin, followed by staining with crystal violet before loading samples on a hemacytometer for counting. To generate adherent haptomonads for imaging, 2 ml of culture at a density of 1 x 10^7^ cells/ml was seeded in a 35 mm poly-L-lysine-coated MatTek dish. The cells were allowed to adhere for 2 hours without rocking. Following 3 washes with BHI, fresh media was added, and dishes were incubated without rocking for 24 hours prior to imaging. To obtain clonal *C. fasciculata* cell lines, serial dilutions of culture were plated on BHI plates made with 0.65% agarose and supplemented with hygromycin (Amresco) [35]. After 4-5 days of growth, parasite colonies were transferred with a pipet tip to a standard liquid culture containing hygromycin.

### Plasmids and cell lines

To obtain cells expressing mitochondrial GFP, the mitoGFP gene was amplified from the pLew100mitoGFP plasmid (a gift from Stephen Hajduk, [36]) using primers 5’-TAT<u>GGTACC</u>tcgtcccgggctgcacg-3’ and 5’-TAT<u>CTCGAG</u>cgcacctccctgctgtgcc-3’ and cloned into pNUSHcH (a gift from Emmanuel Tetaud, [37]) with KpnI and XhoI. The resulting pNUSmitoGFPcH was introduced into *C. fasciculata* CfC1 cells by electroporation (modified from [38]). Briefly, 5 x 10^7^ cells were centrifuged at 1000 rcf for 5 min, washed once with 1 ml electroporation buffer (21 mM HEPES pH 7.4, 137 mM NaCl, 5 mM KCl, 0.7 mM Na_2_PO_4_, 6 mM glucose), and resuspended in 1 ml electroporation buffer. 2.5 x 10^7^ cells were electroporated in a 4 mm gap cuvette containing 40 µg of purified plasmid using a BTX ECM-600 electroporator with a single pulse (475 V, 800 µF, 13 Ω). Cells were allowed to recover for 18-24 hours in BHI supplemented with 10% FBS. Following recovery, hygromycin was added to a final concentration of 80 µg/ml. Transfected cells were passaged 7-12 days later, at which point the hygromycin concentration was increased to 200 µg/ml to select for cells with stronger GFP expression. This cell line was cloned by limiting dilution onto BHI agarose plates to create the cell line CfmitoGFP2.6. A modification of the pNUSmitoGFPcH plasmid containing a ribosomal DNA promoter upstream of the mitoGFP gene was created by amplifying the rDNA promoter from *C. fasciculata* genomic DNA using primers 5’-TATTAT<u>AAGCTT</u>ctgggttatagggggatgct and 5’-TATTAT<u>AAGCTT</u>atcaatcaagtgccgactcc [39, 40]. This fragment was introduced into the HindIII site of pNUSmitoGFPcH to create pNUSrpMitoGFPcH. This plasmid was introduced into the CfC1 line using nucleofection with a Lonza IIb Nucleofector device, the Lonza Human T-cell kit, and program X-001 according to the manufacturer’s protocol. Cells were allowed to recover as described above. The resulting CfrpMitoGFP line (non-clonal) showed comparable fluorescent intensity and pattern as CfmitoGFP2.6, therefore both cell lines were used interchangeably in this study.

### Western blot analysis

To screen clones of CfmitoGFP for maximal expression of the fluorescent protein, protein lysates were created from populations and clones of both the parental CfC1 line and the mitoGFP line. 2 x 10^7^ cells per sample were centrifuged, washed once with PBS, and resuspended in 200 µl of hot Laemmli SDS-PAGE sample buffer. Samples were boiled for 5 min and cleared by centrifugation at 21,000 rcf. 1 x 10^6^ cell equivalents per lane were run on a 12% SDS-PAGE gel. An identical gel was run and Coomassie-stained to confirm equal loading. Resolved proteins were transferred onto PVDF, which was blocked with PBS containing 5% non-fat dry milk and 1% Tween-20 (PBST) and probed with mouse-anti-GFP (Roche) at a dilution of 1:250 in PBST containing 1% milk. The blot was probed with anti-mouse-HRP (Jackson ImmunoResearch) at a dilution of 1:5000, incubated with ECL (BioRad), and imaged with a G:Box gel imager.

### Cell fixation and imaging

To fix cells for fluorescent microscopy, 1 x 10^7^ cells for each sample were centrifuged at 1000 rcf for 5 min and resuspended in 1 ml of BHI. 36% formaldehyde (Sigma-Aldrich) was added to a final concentration of 4% and samples were incubated at room-temperature for 5 min. Fixed cells were centrifuged again and washed twice with 1 ml of PBS. Cells were resuspended in PBS at a density of 1 x 10^7^ cells/ml. 200 µl of this mixture was added to a charged slide (Thermo), which was incubated in a humidified chamber for 10-20 min. Slides were washed once briefly and again for 5 min in PBS in Coplin jars. To permeabilize cells, 200 µl of PBS containing 0.1% Triton X-100 was added to the slides and incubated for 5 min. Slides were then washed as before and stained with 0.2 µg/ml DAPI in PBS for 5 min. Slides were washed again and mounted in 90% glycerol in PBS. Wide-field fluorescent imaging was performed on a Zeiss Axioscope.A1 upright LED fluorescence microscope equipped with a Zeiss AxioCam ICm1 camera. Confocal imaging of fixed cells was performed on a Leica SP5-II confocal microscope. Length and width of cells was measured in ImageJ. To measure mitochondrial area, maximum projections of confocal Z-stacks were generated in ImageJ. After applying a fluorescence threshold, the area was calculated. Nodes were counted manually in ImageJ max projections. Statistical analyses were performed using GraphPad Prism. Data from individual samples was first analyzed for normality using the D’Agostino & Pearson normality test. If all samples being analyzed showed a normal distribution, an ANOVA multivariable analysis or an unpaired t-test was used to test for significance. If at least one sample did not show a normal distribution, a Mann-Whitney test was used.

### Synchronization

Synchronization was performed as in [41]. Briefly, 2.5 mM (final) hydroxyurea (HU, Sigma) was added to a 20 ml culture at a density of 1 x 10^7^ cells/ml. After six hours of treatment, cells were centrifuged, washed once with 20 ml BHI without HU and then resuspended in fresh BHI. A portion of culture was immediately removed and processed for fixed cell microscopy as described above (0 hour). Additional samples were removed and processed every 30 min until 150 min. A sample was also prepared from an asynchronous culture at comparable density. For each sample, two slides were made. One set of slides were analyzed by wide-field fluorescent microscopy to assess the efficiency of the synchronization procedure, while the other set was analyzed by confocal microscopy. For fluorescence microscopy-based quantitation the mean and standard error of three independent replicates is shown except for times 0 and 0.5 h post-release which were examined in two replicates.

### Treatment with mdivi-1

Mdivi-1 (Sigma) was resuspended in DMSO to a final concentration of 20 µg/ml. Addition of this solution to cell cultures produced a slight precipitate. This precipitate did not seem to affect the reproducibility of our results, but our effective concentrations of mdivi-1 may be lower than the indicated amount. DMSO was added to one culture to serve as a vehicle control. At the times indicated in the figure, a sample of 1 x 10^7^ cells was removed and prepared for fluorescence microscopy as described above. Given the processing time, the minimum time of treatment was 5 min. For experiments using MitoTracker Red CMXRos (Thermo), cells were preincubated with either DMSO or 100 µg/ml mdivi-1 for 30 minutes at 27 °C. Cells were spun at 1000 rcf for 5 min, supernatants removed, and pellets resuspended in BHI media containing 0.2 µM MitoTracker Red and pre-warmed to 27 °C. Mdivi-1 or DMSO was then added back to its initial concentration and the cells were incubated for a further 30 min at 27 °C prior to fixation and preparation for microscopy as described above. Alignments and percent identities between dynamin related proteins from *C. fasciculata, S. cerevisiae*, and *H. sapiens* were performed in ClustalOmega [42]. Accession numbers for protein sequences were CfDLP: CFAC1_200029900; ScDNM1: NP_013100.1; HsDRP1: NP_036192.2.

### Live cell imaging

To immobilize swimming nectomonads for live cell imaging two methods were used. First, an agarose pad was prepared by adding 100 µL of BHI containing 1% low melting point agarose (IBI Scientific). This solution was boiled, equilibrated to 65 °C, then added to a glass slide. Another slide was placed on top. After 10 min the top slide was gently removed and 20 µl of a mid-log phase culture (∼3 x 10^7^ cells/ml) was added to the center of the agarose pad. A coverslip was added and sealed with VaLaP. In the second method, a BHI-agarose mixture was prepared as before, equilibrated to 65 °C, then mixed 1:1 with a culture of swimming cells. 20 µl of this mixture was added to a slide, cover-slipped, and sealed with VaLaP. Both methods were reasonably effective at immobilizing living *C. fasciculata* parasites. Cells with motile flagella (an indication of cell viability) but an immotile cell body were selected for imaging. For live-cell imaging of adherent haptomonads, cultures were prepared as described above. Cells were imaged with either a Yokagawa CSU X-1 spinning disk confocal head configured on an inverted Leica DMI4000 microscope with a Hamamatsu EM512 camera or a Leica SP5 II laser confocal microscope on a DMI6000 stand as indicated. On the spinning disk confocal Z-stacks were taken every 2 min for up to 60 min without significant photobleaching or phototoxicity. On the laser scanning confocal Z-stacks were taken every 12-30 seconds for up to 40 min and a resonant scanner was used for rapid imaging. Both confocal systems were outfitted with environmental controls which were set to 27 °C during imaging. Some time-lapse confocal images were processed with Huygens deconvolution software (Scientific Volume Imaging).

## Results

### Mitochondrial GFP is a marker for the *C. fasciculata* mitochondrion

To image the *C. fasciculata* mitochondrion, we obtained a mitochondrial GFP (mitoGFP) construct in which a 14-amino acid mitochondrial targeting sequence from *T. brucei* is fused to GFP [36], targeting the protein to the mitochondrial matrix. The activity-based dye MitoTracker Red CMXRos also labels the mitochondrion, but in *C. fasciculata* tends to accumulate in one or two bright puncta which saturate before the rest of the organelle shows robust signal. We therefore created a drug-selectable mitoGFP expression construct for *C. fasciculata* [37]. After introducing this construct into wild-type cells, we observed a fluorescence pattern consistent with kinetoplastid mitochondria and which colocalized with MitoTracker Red (Fig 1). Expression of mitoGFP is heterogeneous within the cell population, as one would expect for an episomal construct (Fig 1A, [37]). By screening clones of this cell line, we were able to select a clone with a higher overall signal, presumably because it maintains more copies of the expression construct (S1 Fig). In parallel studies we modified our expression construct by introducing an rDNA promoter upstream of the mitoGFP gene. This resulted in a strong, yet still heterogeneous, signal that also colocalized with MitoTracker Red. Both cell lines are used in this study as they showed an identical pattern and were of comparable brightness. We saw no association between the strength of the mitoGFP signal and the stage of the cell cycle.

**Fig 1.**
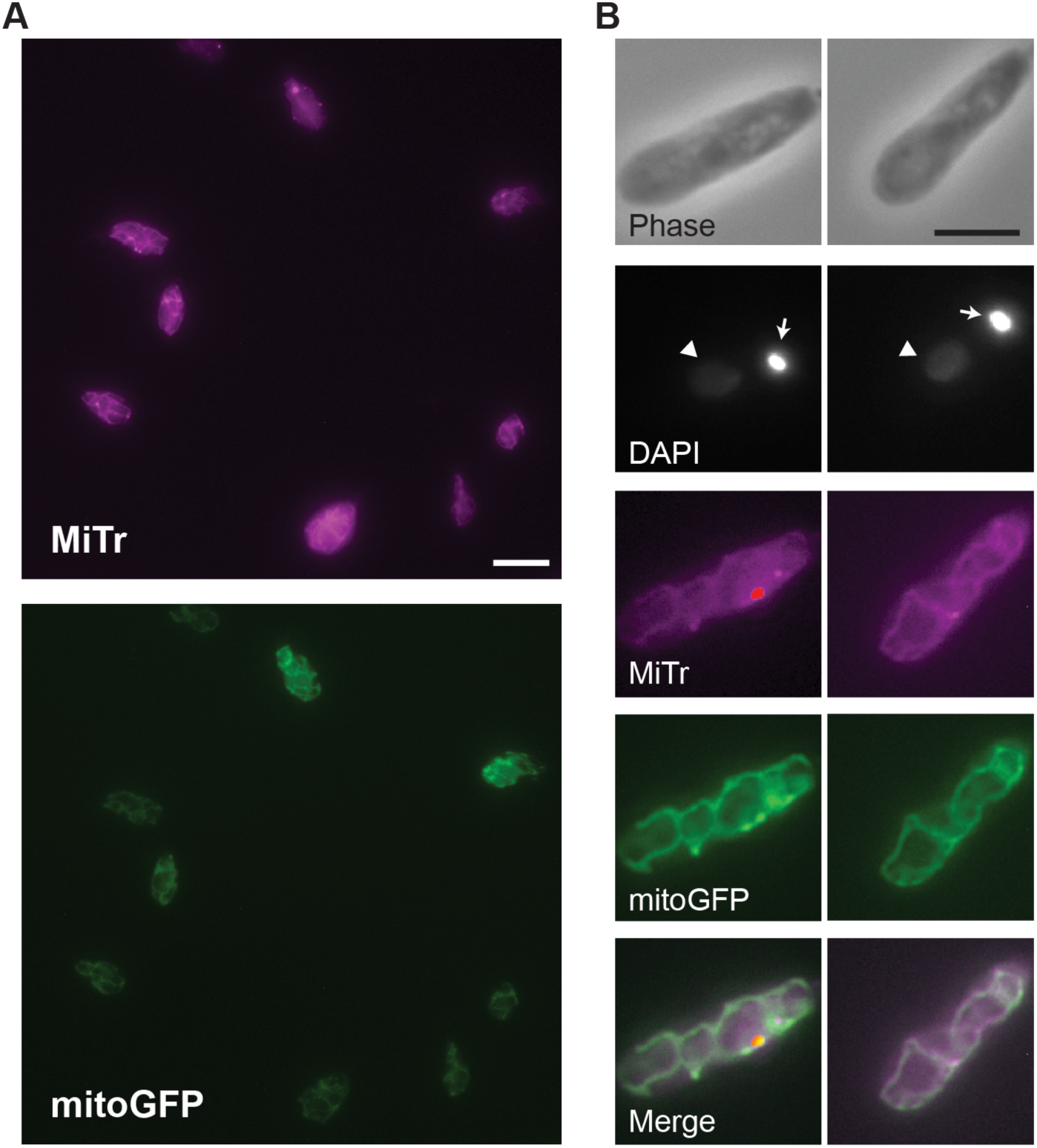
mitoGFP is a mitochondrial marker in *C. fasciculata*. A) Mitochondrial staining by the potential-dependent dye MitoTracker Red (MiTr, top panel) and by mitochondrial-targeted GFP (mitoGFP, lower panel). mitoGFP expression is driven by an ectopic plasmid causing heterogeneous fluorescence among cells. Scale bar is 10 µm. B) Higher magnification images of two representative cells show colocalization between MiTr and mitoGFP signals. The phase contrast shows the cell boundaries and nuclear DNA (arrowhead) and kDNA (arrow) are stained with DAPI. In merged images MiTr is shown in magenta and mitoGFP in green. Scale bar is 5 µm.

### Mitochondrial biogenesis during the *C. fasciculata* cell cycle

We first examined the overall shape of the mitochondrion in logarithmically-growing *C. fasciculata* cells. Consistent with what has been reported for other kinetoplastid parasites, we found that the mitochondrion takes the shape of a branched tubular network that extends throughout the cell (Fig 2A). In many cells, there was a clear space with fewer mitochondrial tubules in the part of the cell that contains the nucleus. We also noted that many cells contained areas of thickened mitochondrial tubules. This variability in mitochondrial diameter did not appear to associate with any particular phase of the cell cycle.

**Fig 2.**
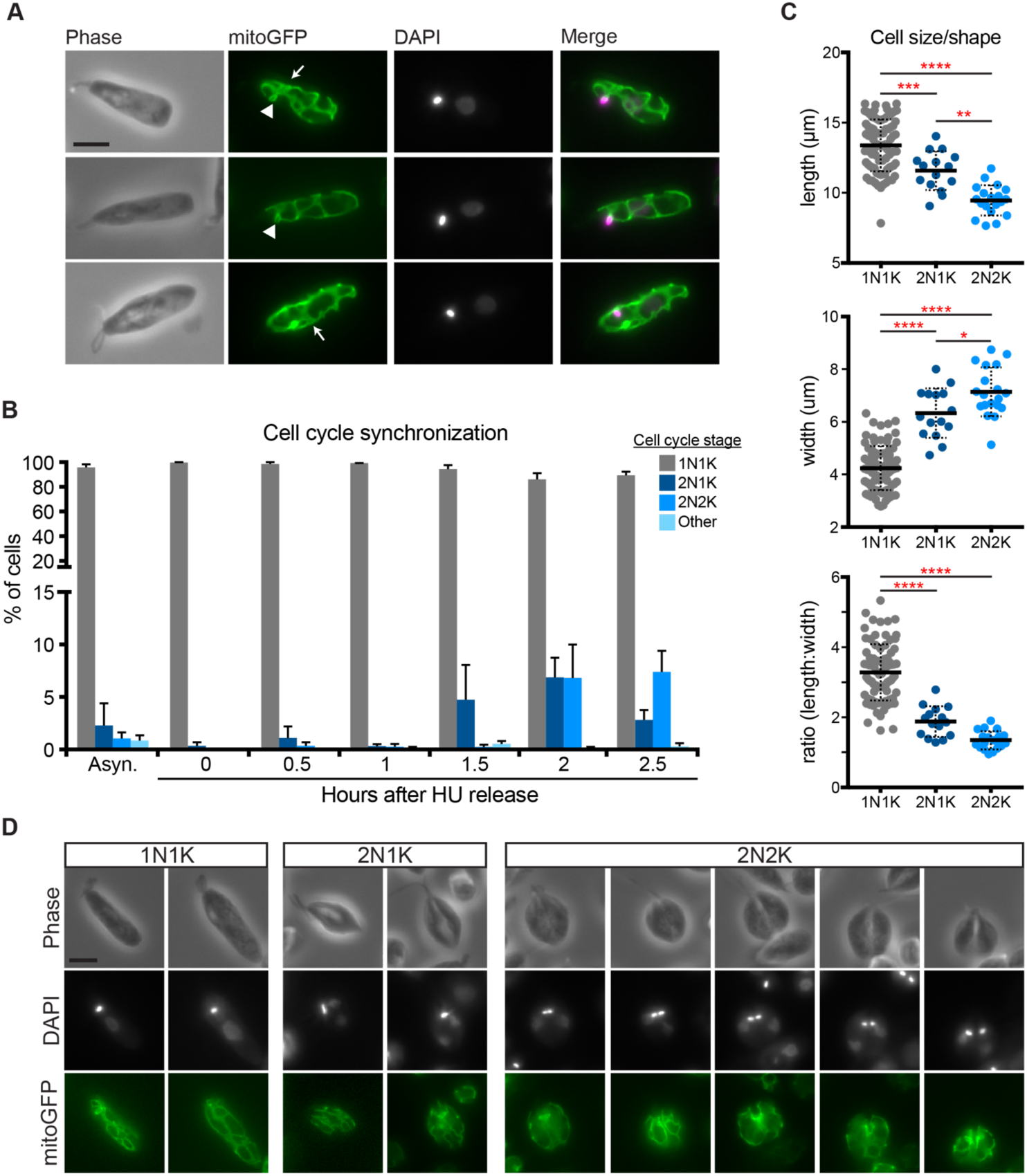
Mitochondrial shape during the cell cycle of *C. fasciculata*. A) Typical mitochondrial morphology in 1N1K (G1 phase) cells as shown by mitoGFP. Arrowheads show a small mitochondrial loop that indicates the presence of the kDNA disk. Arrows point to areas containing thickened mitochondrial tubules. DAPI signal is shown in magenta in merged images. B) Quantitation of number of cells in different phases of the cell cycle in asynchronous cultures and in hydroxyurea (HU)-synchronized cultures at various times after removal of HU. This procedure allows for enrichment of mitotic cell cycle phases and demonstrates cell cycle progression from 1N1K to 2N1K, 2N2K, and back to 1N1K. N, nucleus. K, kDNA. C) Synchronized cells (n=119) in different phases of the cell cycle according to DAPI staining were measured. P-values indicated *<0.05, **<0.005, ***=0.001, and ****<0.0001 by ANOVA. D) Examples of typical mitochondrial shapes found in 1N1K, 2N1K, and 2N2K cells. Note that the nuclei are not symmetrically positioned with respect to the cell body in 2N1K cells. The last three columns show cells undergoing cytokinesis.

Kinetoplastids, like *C. fasciculata*, have only a single mitochondrion. In these cells, mitochondrial biogenesis and division must be timed with the cell cycle. In addition, each mitochondrion contains a single mitochondrial nucleoid, called the kinetoplast DNA (kDNA), which consists of thousands of catenated DNA circles. It is well-known that replication and division of the kDNA also occurs within a particular cell cycle stage [43–46]. To describe the mitochondrial duplication and division cycle of *C. fasciculata*, we first examined asynchronous cultures of mitoGFP-expressing cells. Staining with DAPI and counting the number of nuclei and kDNA in each cell allows for a rough approximation of cell cycle stage. In *Crithidia*, unlike in *T. brucei*, the nucleus (N) divides first, followed by the kDNA (K). This is also the order of events in the *Leishmania* cell cycle [45]. Therefore, cells progress from 1N1K to 2N1K and 2N2K before undergoing cytokinesis and returning to 1N1K (Fig 2B). In asynchronous culture, more than 90% of the cell population is 1N1K, indicating that the stages following mitosis and kDNA division are brief (Fig 2B). To enrich for these stages, we synchronized our cultures using hydroxyurea (HU) [41, 44]. This nucleoside analog causes cells to arrest and accumulate in G1 (1N1K). Removal of the drug releases the block, allowing cells to progress through the cell cycle. Using our synchronization enrichment protocol, we were able to observe ∼11% 2N1K cells at 1.5 h post-release and ∼12% 2N2K cells at 2 h post-release (Fig 2B). This is compared to approximately 1% in asynchronous cultures.

When we analyzed mitoGFP cells by fluorescence microscopy, 2N1K and 2N2K cells appeared wider and shorter than 1N1K cells. This is similar to the pattern of cell cycle progression in *Leishmania mexicana* [45]. To quantitate this, we measured the length and width of cells in different phases of the cell cycle in our partially synchronized cultures. Indeed, as cells progressed from 1N1K to 2N2K their average length decreased while their width increased (Fig 2C). This resulted in a morphologically symmetrical cell that can be divided almost exactly in half during cytokinesis, followed by cell elongation in G1. However, when we examined the pattern of DAPI staining in 2N1K cells we observed that the positioning of the nucleus was not perfectly symmetrical (Fig 2D). Again, this is consistent with what has been observed in *L. mexicana* [45]. This asymmetry resolves as cells progress to the 2N2K stage, when the kDNA is positioned symmetrically over the cell division plane.

The cell cleavage furrow begins, as in other kinetoplastids [47], at the anterior end of the cell between the two flagella and moves posteriorly until the cells separate. During this process, the mitochondrion remains branched and appears to be equally partitioned between daughter cells (Fig 2D). Some mitochondrial tubules are found close to the cell periphery. In 2N2K cells, these tubules often formed a “V” shape, where the point of the “V” coincided with the progressing cytokinesis furrow (Fig 2D).

### Distinct mitochondrial morphologies in replicating *C. fasciculata*

To perform a more detailed analysis of mitochondrial morphology during the cell cycle, we used confocal microscopy to image mitoGFP cells following synchronization. We took confocal z-stacks through 83 1N1K cells, 15 2N1K cells, and 16 2N2K cells from two separate experiments. Again, we observed that the majority of mitochondrial networks (∼81%) were mixtures of thin tubules and thicker, flatter tubules, while the remaining cells contained mitochondrial tubules of fairly uniform diameter.

This analysis also revealed some cell cycle stage-specific mitochondrial morphologies (Fig 3A, B). In 43% of G1 phase cells (1N1K), the mitochondrion is a network of closed loops. In the remaining cells, there were free mitochondrial ends extending towards the anterior end of the cell. These open networks were commonly found in wider cells with enlarged or mitotic nuclei. Consistent with this, in 2N1K cells there were a greater number of cells with open mitochondrial networks (87% in 2N1K compared to 57% in 1N1K). Closed networks predominate again during the 2N2K stage, with only 13% of 2N2K cells having open mitochondrial networks.

**Fig 3.**
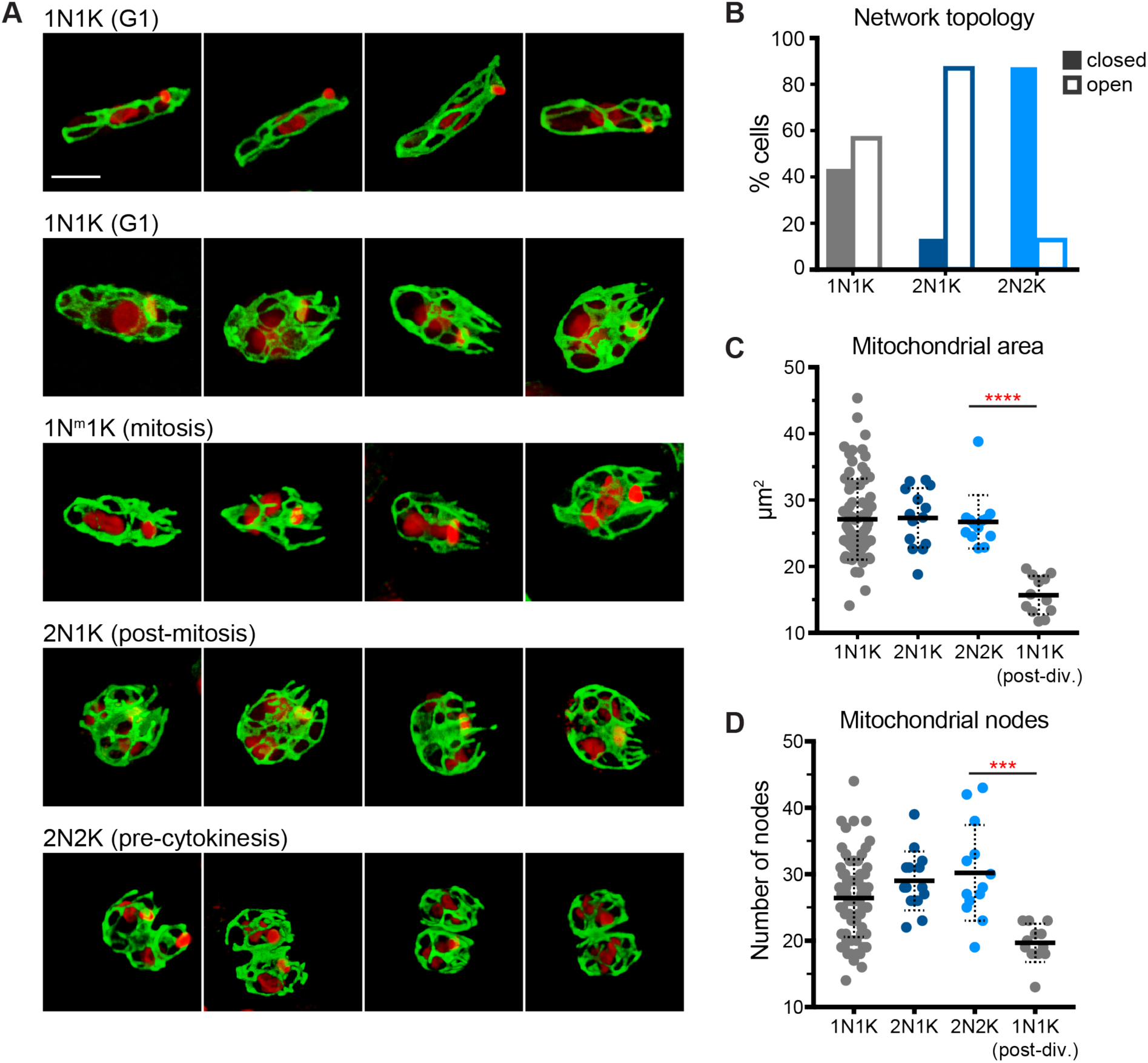
Confocal analysis of mitochondrial shape during the cell cycle of *C. fasciculata*. A) Examples of maximum projections of fixed, synchronized *C. fasciculata* mitoGFP-expressing cells as imaged by confocal microscopy. Mitochondria are shown in green and DAPI in red. Cell cycle stages are shown above each row. The first row shows all “closed” networks. Rows 2-4 show networks that are “open” at the anterior end (containing tubule end points rather than loops). Each image is oriented such that the anterior of the cell is on the right. Scale bar is 5 µm. B) Quantitation of the percent of cells (n=114) in each cell cycle stage that have closed or open mitochondrial networks at their anterior end. C) Mitochondrial area in z-stack maximum projections as a function of cell cycle stage (n=115). **** indicates P-value <0.0001 by Mann-Whitney test. D) The numbers of mitochondrial nodes, or branch points, were counted manually in maximum projections (n=115) *** indicates P-value of 0.0002 by unpaired t-test with Welch’s correction.

To examine mitochondrial biogenesis and topology during the cell cycle we analyzed maximum projections of each z-stack to quantitate mitochondrial area and number of network branch points (Fig 3C and D). A branch point was defined as a position at which two or more mitochondrial tubules diverged. Interestingly, we found a wide range of mitochondrial areas in 1N1K cells compared to more narrow ranges for 2N1K and 2N2K cells. The mean mitochondrial area in each of these categories is approximately the same. When we separated 1N1K cells that had clearly just divided or were minimally connected to another 1N1K cell by a narrow cytoplasmic bridge and treated them as a separate category, we found that their mean mitochondrial area was approximately half that of the other cell types. This implies that the mitochondrion is divided in half during cell division, and then grows to its full size during G1. This would at least partially explain the variability in mitochondrial size that we observe in our 1N1K population. Some 1N1K mitochondria are actually larger than those found in 2N1K and 2N2K cells, indicating that, during G1 growth, the mitochondrion might actually overshoot its target size before returning to a steady-state prior to division. This additional mitochondrial biogenesis may be necessary to support rapid cell growth during this cell cycle stage. By the 2N1K stage, the mitochondrion has returned to its normal size, which is similar in all 2N1K cells analyzed. Similar to mitochondrial area, the number of mitochondrial branch points remained relatively constant except in recently divided cells, when it was halved. These data suggest that both cell and mitochondrial growth occur primarily in G1, and that the size of the mitochondrion tracks closely with cell size.

### Treatment with the inhibitor mdivi-1 does not affect mitochondrial division

In *T. brucei*, knockdown of the dynamin-like protein TbDLP caused inhibition of mitochondrial division as well as cytokinesis through an unknown checkpoint connecting division of the organelle to division of the entire cell [30]. Such a checkpoint would ensure mitochondrial inheritance in an organism with only one mitochondrion per cell. We identified a single DLP in the *C. fasciculata* genome. To determine if CfDLP is required for mitochondrial division in *C. fasciculata*, we sought to block its activity. Since RNAi is not yet established in these organisms and CfDLP is expected to be essential, we used a chemical inhibitor of dynamin-like proteins called mdivi-1. It has been reported that mdivi-1 interferes with the activity of dynamin-like proteins (but not classical dynamins) in both yeast and human cells [48]. We thought that mdivi-1 would also block CfDLP given the conservation between CfDLP and its yeast and human orthologs (40% identity between CfDLP and either the human or the yeast protein). Surprisingly, we found that treatment with mdivi-1 had only a minor effect on cell growth (Fig 4A) and no discernable effect on the shape of the mitochondrial network as assessed by mitoGFP (Fig 4B). Unexpectedly, treated cells appeared reduced in size compared to untreated cells or cells treated with a vehicle control (Fig 4B). To quantitate this effect, we measured cell lengths during a time-course of mdivi-1 treatment (Fig 4C). To eliminate effects of cell cycle on size, we measured only 1N1K cells, although 2N1K and 2N2K cells also appeared smaller. This analysis confirmed the effect of mdivi-1 on cell size and showed that the effect occurred very rapidly, being already maximal (1.4-fold reduced) after 40 min after treatment. In fact, the effect occurs within as little as 5 minutes, which is as quickly as we can process the cells (Fig 4D).

**Fig 4.**
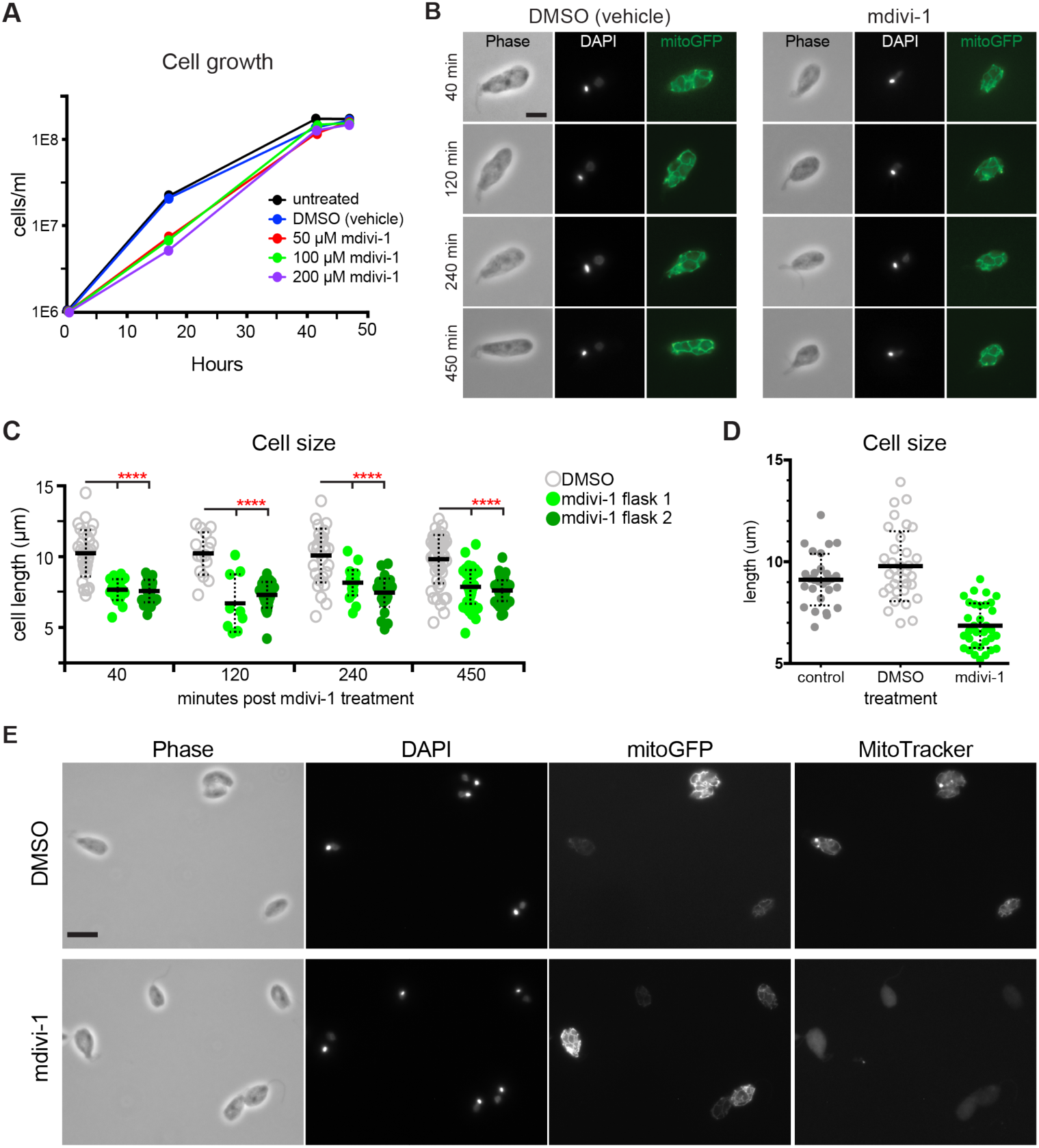
Treatment with mdivi-1 does not inhibit mitochondrial division in *C. fasciculata*. A) Growth curve of *C. fasciculata* mitoGFP-expressing cells in the presence of different concentrations of mdivi-1 or vehicle control (DMSO) equivalent to that of the 200 µM concentration of mdivi-1. B) Representative examples of cells treated with vehicle (DMSO) or 100 µM mdivi-1 for the times indicated. Cells were fixed and stained with DAPI. Mitochondrial shape is visualized with mitochondrial GFP (mitoGFP). Scale bar is 5 µm. C) Measurements of cell length in DMSO or mdivi-1-treated 1N1K cells (two separate flasks). 40 min, n=78; 120 min, n=55; 240 min, n=84; 450 min, n=165. The difference between DMSO-treated and each mdivi-1-treated flask was significant for all time points (****, P<0.0001, unpaired t-test). D) Measurements of cell length in untreated, DMSO (vehicle), and 100 µM mdivi-1 (n=95). Time of treatment was approximately 5 min. Only 1N1K cells were measured. (*****, P<0.0001, unpaired t-test). E) mitoGFP-expressing cells were exposed to either DMSO or 100 µM mdivi-1 for 30 min followed by MitoTracker Red (MiTr) staining for 30 min. Cells were fixed and stained with DAPI. Representative images are shown. Scale bar is 10 µm.

Given how quickly the cells respond to the inhibitor, we decided to use MitoTracker Red CMXRos to determine if mdivi-1 affects mitochondrial membrane potential in *C. fasciculata*. In control, vehicle-treated cells, the MitoTracker signal accumulates in the mitochondrial tubules (Fig 4E). As indicated previously, bright puncta, typically localized at the anterior of the cells, were observed. In contrast, in mdivi-1-treated cells the MitoTracker dye did not accumulate in the mitochondrion. Instead, the dye was distributed throughout the cytosol (Fig 4E). The mitoGFP signal was present, indicating that the organelle is intact and that mdivi-1 indeed disrupts mitochondrial membrane potential. The speed at which we observe a phenotype, and the loss of membrane potential, means that mdivi-1 is probably affecting something other than CfDLP. For example, some of the effects of mdivi-1 have been attributed to inhibition of mitochondrial respiratory activity at complex I [49]. The function of CfDLP will probably remain undetermined until methods for inducible protein knockdown are established in *C. fasciculata*.

### The *C. fasciculata* mitochondrion is a dynamic network

To address whether mitochondrial biogenesis in G1 phase cells requires events resembling mitochondrial dynamics, we imaged mitochondria in live *C. fasciculata*. For swimming nectomonads, we immobilized asynchronous mitoGFP parasites and imaged them on a confocal microscope (Fig 5). We observed that mitochondrial networks are dynamic even in G1 phase cells (those with a single flagellum, Fig 5A and S1 and S2 Movies). The topology of the network was constantly changing as tubules divided, fused, and emerged from existing tubules. Branch points were also observed to migrate along tubules. In some instances, complex net-like structures resembling fenestrated sheets of mitochondrion would appear transiently before resolving into tubules and loops. The position of the kDNA within the mitochondrial network was usually visible as a dark area surrounded by a small loop of fluorescence signal (Fig 5A). This has also been observed in *T. brucei* and was named the “kDNA pocket” [50]. The kDNA serves as a landmark, allowing us to ensure that dynamic events are not due to the cell rolling or shifting on the slide. We manually scored remodeling events in 17 cells to determine their frequency (Fig 5B). Though some cells were assayed for longer, we calculated the frequency of events during a 30-minute interval because photobleaching was negligible over this period. In total, the average length of time between network remodeling events of any category was 3.1 minutes, or 12.7 events per 30 minutes of observation. The frequency and types of events recorded for *C. fasciculata* are consistent with what has been reported for yeast [51] and mammalian [52] mitochondrial networks and implies that this phenomenon may be widely conserved in eukaryotic cells. We found that tubule fusions were the most common event observed, with an average of 5.8 events in a 30-minute period. These fusion events included both side-to-side tubule fusions and end-to-side fusions. Over this same time period, fission, or division of a tubule, was observed an average of 2.5 times, while budding of a new tubule from an existing one was observed 1.5 times. Branch point sliding occurred with a greater frequency, with 2.8 events per 30 minutes, while the rarest event was the appearance of a fenestrated sheet, which was only observed in 6 of the 17 cells imaged and in those only once per observation period.

**Fig 5.**
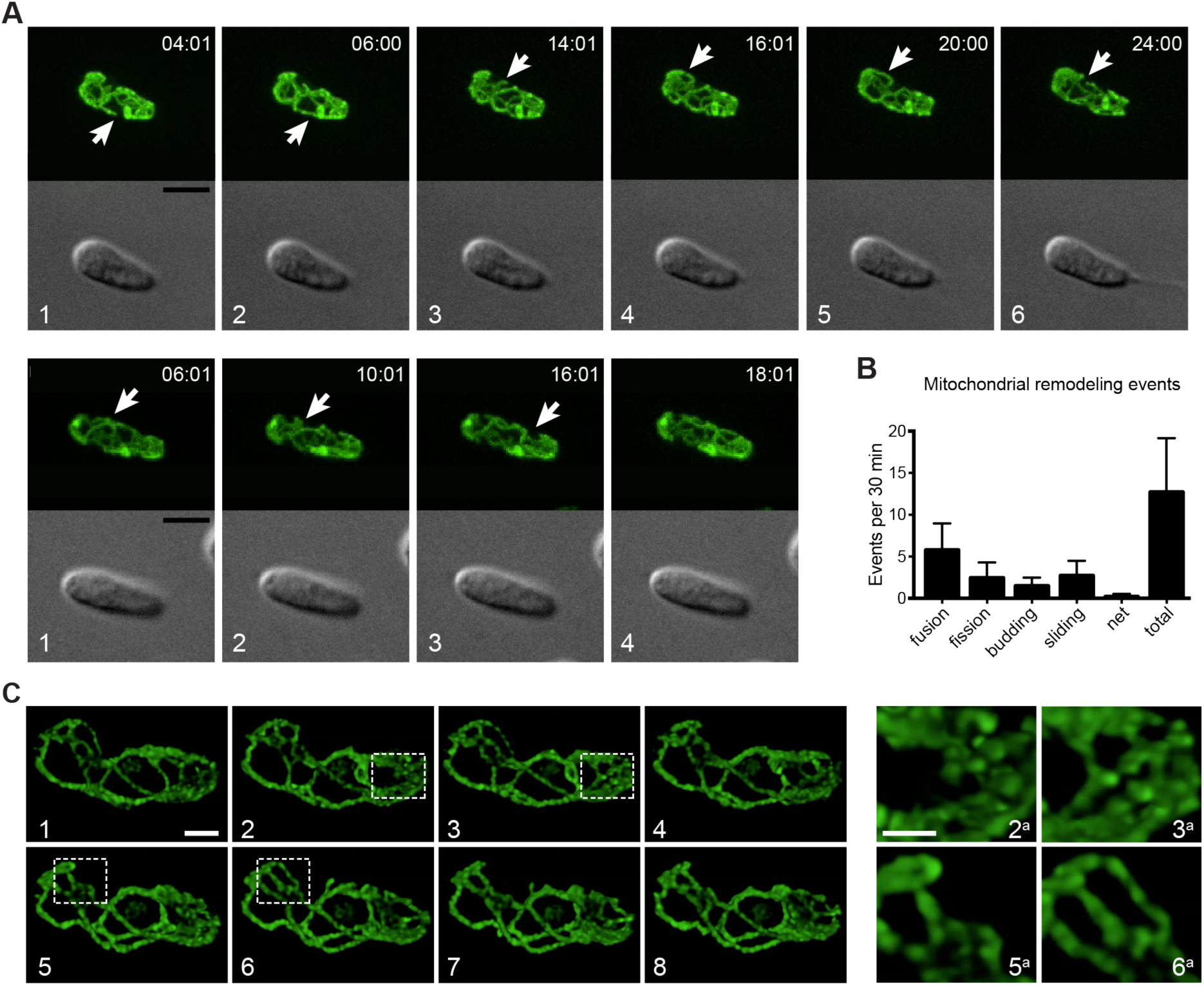
Mitochondrial dynamics in *C. fasciculata*. A) Frames from live-cell imaging of two cells by spinning-disk confocal microscopy. Cells are expressing mitoGFP. Mitochondrial remodeling reactions are indicated by arrows. Times are given as h:min:s. White numbers indicate frame sequence. Not all frames in the time-course are shown (see S1 and S2 Movies) B) Quantitation of mitochondrial remodeling events in spinning disk confocal experiments. Z-stacks of 17 cells were generated every 2 min for an average of 49 min. The resulting movies were then analyzed for different types of mitochondrial remodeling events. The frequency of each event per minute was calculated then multiplied by 30 to give number of events/30 min time period. The black bar (total) indicates the sum of all the different types of events shown by the colored bars. C) Live-cell imaging of mitochondrial remodeling with a laser-scanning confocal microscope and deconvolution. White numbers indicate different frames. Boxes indicate areas in which remodeling occurs between frames. These areas are shown enlarged (with corresponding numbers) in the right panels.

We also visualized these dynamic events using laser scanning confocal microscopy, enabling the use of deconvolution to view mitochondrial network remodeling in more detail (Fig 5C). This analysis revealed that some regions of the network were more dynamic than others, with rapid remodeling (mainly fusions, fissions, and branch point migration) occurring at the anterior and posterior ends of the cell, and less frequent remodeling in the center of the cell, where such events might be physically restricted by the presence of the nucleus.

### Mitochondrial dynamics in adherent, non-motile *C. fasciculata*

An advantage of working with *C. fasciculata* is that cultured cells will readily differentiate into a naturally adherent, non-motile stage [34]. If culture flasks are agitated with gentle shaking, the majority of cells will remain in the swimming or nectomonad form, which is highly motile with a full-length flagellum. In stationary cultures, a fraction of cells will adhere to the tissue culture plastic via their flagellum and differentiate into the non-motile haptomonad form. Haptomonds can grow and divide while adhered to a plastic or glass surface. The flagellum in haptomonads is dramatically shorter, and usually does not emerge from the cellular invagination called the flagellar pocket. We observed live mitoGFP-expressing haptomonads by confocal microscopy and found that their mitochondrion was a branched network similar to that of nectomonads. In addition, remodeling events consistent with mitochondrial dynamics were readily apparent (Fig 6A and S3 and S4 Movies). In particular, the area of the mitochondrion containing the kDNA was very kinetic, appearing as a loop of mitochondrial matrix at the end of a mitochondrial tubule stalk. The movement of the kDNA disk may be due to motility of the shortened haptomonad flagellum, or through another, unknown mechanism. We observed similar classes of remodeling events as were seen in nectomonads, including the transient appearance of fenestrated sheets of mitochondrion, which quickly resolved into tubules (Fig 6B and S4 Movie).

**Fig 6.**
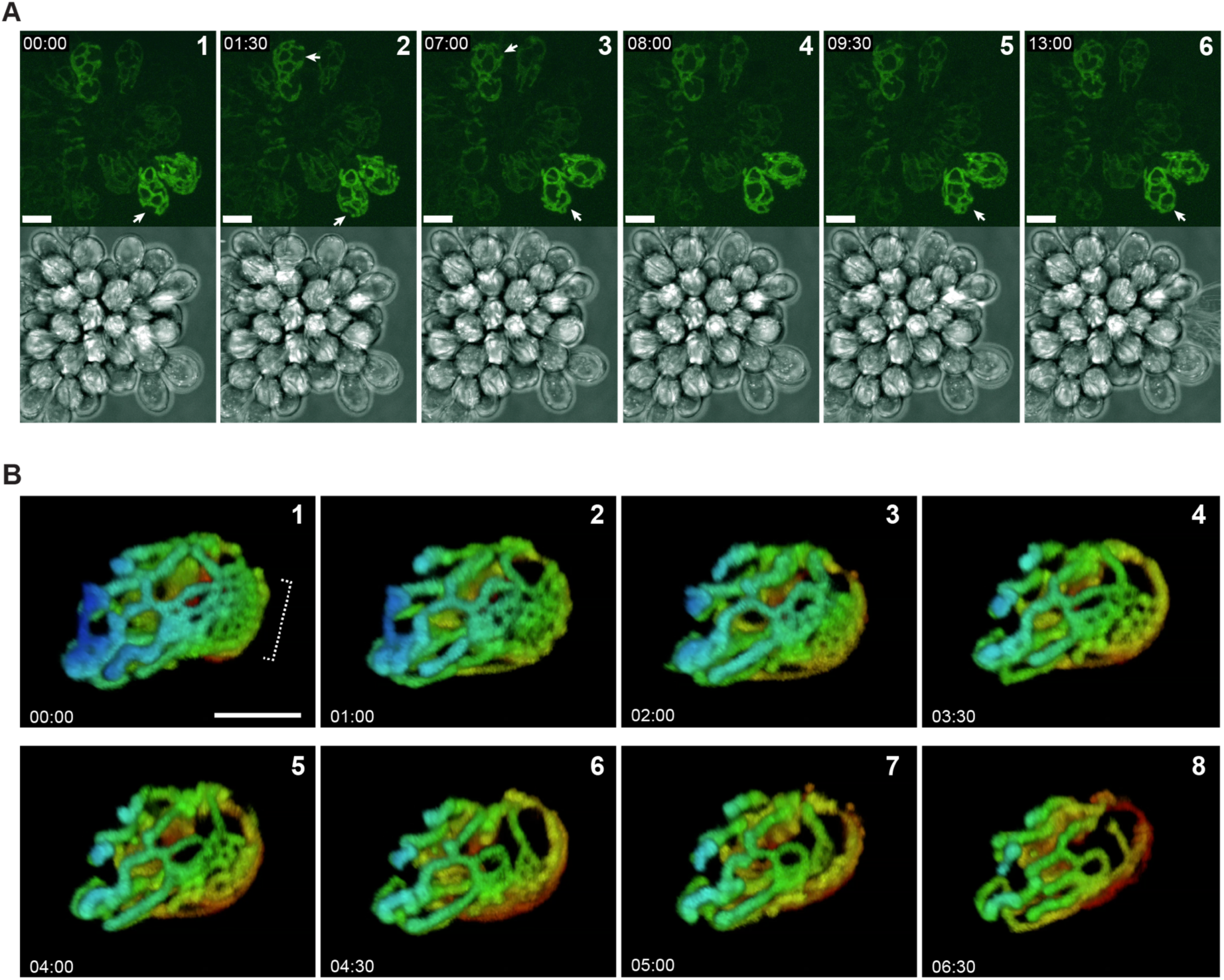
Adherent *C. fasciculata* haptomonads also show rapid mitochondrial remodeling. A) Max projections of a typical rosette of *C. fasciculata* imaged using a laser scanning confocal microscope. Mitochondria in live cells were visualized using mitoGFP. Mitochondrial remodeling reactions are indicated by arrows. Times are given as mm:ss. White numbers indicate the sequence of frames. Note that not every frame is shown (see S3 Movie). Scale bar is 10 µm. B) Time course of mitochondrial remodeling in a single adherent cell. Each frame is a max projection that has been color-coded by depth. White numbers indicate the sequence of frames. Note that not every frame is shown (see S4 Movie). Bracket indicates the formation of a net-like structure that then is rapidly remodeled to tubules during the course of our observation. Scale bar is 5 µm.

### The coordination of kDNA division, mitochondrial division, and cytokinesis

Our fixed cell studies indicate a tight coupling between mitochondrial division and cytokinesis. To explore this relationship further, we visualized these processes in live cells. Again, we observed that mitochondrial division occurs shortly before cytokinesis in both nectomonad and haptomonad cells (Fig 7A and B and S5-S7 Movies). The progress of mitochondrial division matched that of the cleavage furrow, beginning at the anterior end of the cell (between the two daughter flagellar pockets) and progressing posteriorly. The rest of the mitochondrion continued to undergo remodeling during the 2N1K and 2N2K stages, indicating that mitochondrial dynamics is not limited to a particular cell cycle stage (S7 Movie). However, we noted increased fusion of tubules at the midline of the cell at the position where the cytokinesis plane would pass through. This might ensure the even mixing and distribution of mitochondrial components prior to division.

**Fig 7.**
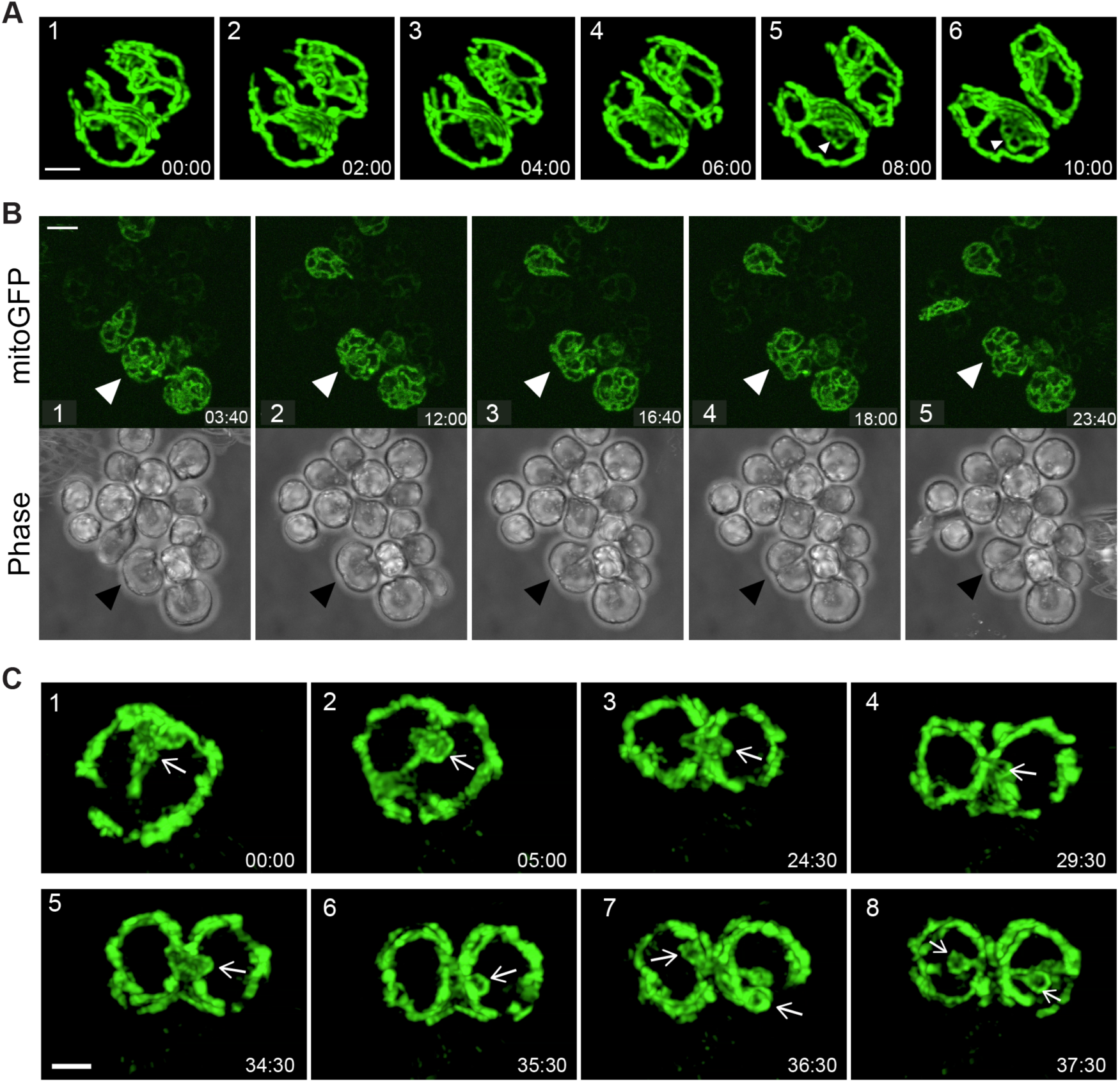
Mitochondrial division occurs just prior to cytokinesis in *C. fasciculata*. A) Time-lapse of mitochondrial/cell division in a swimming nectomonad immobilized in low melting point agarose. Image is a max projection of a Z-stack generated on a spinning disk confocal microscope and subjected to deconvolution. Times are given in mm:ss and numbers indicate the sequence of frames. The anterior of the cell is on the left side of each image. Complete mitochondrial division seems to occur between frames 3 and 4. A fenestrated sheet appears in the lower cell in frames 5 and 6 (arrowheads). Not all frames are shown (see S5 Movie). Scale bar is 2 µm. B) Mitochondrial and cell division of adherent haptomonads. Arrowheads indicate a dividing cell. Times are given in mm:ss and numbers indicate the sequence of frames. Not all frames are shown (see S6 Movie). Scale bar is 10 µm. C) Time lapse of a dividing haptomonad. The presumed position of the kinetoplast (identified by the small loop of mitochondrial matrix surrounding it, is indicated by white arrows. Numbers indicate the sequence of frames. Not all frames are shown (see S8 Movie). kDNA division occurs in frame 6-7. Scale bar is 2 µm.

Our live cell imaging studies indicate that division of the kDNA is also temporally linked to cytokinesis. Prior to the initiation of cytokinesis, the kDNA moves from the anterior end to the center of the cell, straddling the division plane (Fig 7C and S8 and S9 Movies). The kDNA pocket appears as a single dark spot surrounded by a mitochondrial loop. This structure is larger than that in G1 cells since the kDNA has already replicated by this point. Shortly after the appearance of the cytokinesis furrow, the kDNA is divided in half and two smaller loops are visible (Fig 7C and S8 and S9 Movies). The process of kDNA division is rapid, occurring in under 10 minutes. This is interesting since the exact mechanism for dividing this complex structure is not understood. In *C. fasciculata*, kDNA minicircles are removed from the center of the network but are reattached evenly along the entire periphery of the disk [53]. This is facilitated by relative movement of the kDNA disk between the two polar sites where reattachment occurs. It is not known what process drives this movement, or how it is accommodated by the system of filaments that link the kDNA to the basal body [54]. The spinning kDNA network enlarges during replication, and retains its disk shape, after which it is divided precisely down the middle. To our knowledge, this is the first time this process has been observed in real time.

## Discussion

Here we present evidence for rapid mitochondrial dynamics in the monoxenous kinetoplastid parasite, *C. fasciculata*. These events occur in both motile and non-motile cells and in all stages of the cell cycle. Mitochondrial remodeling reactions may be important for maintaining the size and topology of the mitochondrial network, and for allowing its growth and division during the cell cycle. We have shown that the majority of both cell growth and mitochondrial biogenesis occurs in G1, and that the morphogenic events of the *C. fasciculata* cell cycle resemble those in *Leishmania*. Following mitosis, the *C. fasciculata* mitochondrion is divided in half in a process that is coincident with cytokinesis. This division process is not impaired by treatment with the putative dynamin inhibitor mdivi-1 which instead reduced cell size and mitochondrial membrane potential. We have also shown the spatial and temporal coordination of kDNA division with the cell cycle through the examination of live cells.

The single mitochondrion of kinetoplastid parasites has been presumed to be a relatively static network, with significant remodeling limited to division of the organelle prior to cell division [30] and differentiation between life cycle stages (in the case of *T. brucei* [55, 56]). In all kinetoplastids, mitochondrial fission is required for division of one organelle into two prior to or during cytokinesis. In *T. brucei*, this activity is mediated by TbDLP, which also plays a role in endocytosis [30]. There have been several studies on the two nearly identical TbDLPs, yet none of them show a clear effect on mitochondrial shape or dynamics such as that produced by depletion of the orthologous protein in yeast or mammalian cells [28–30]. The fact that knockdown of the only proposed mitochondrial fission activity in *T. brucei* has little effect on basic network structure implies that mitochondrial shape is maintained by other mechanisms. Without a “typical” role for DLP, kinetoplastids may serve as an excellent platform to discover dynamin-independent mechanisms of mitochondrial remodeling, which may be conserved in other organisms. A recent report on the dynamics of mitosomes in *Giardia intestinalis* parasites found that they undergo fission but not fusion, that fission is coordinated with the cell cycle, and that it occurs independently of dynamin-like proteins [57].

We have observed mitochondrial tubule fission events in *C. fasciculata* that occur continuously during the cell cycle. Whether these reactions are mediated by CfDLP is an open question, and answering it will require a genetic approach. We have also detected the dynamic formation of mitochondrial branches and the creation of fenestrated sheets. Branching may be important for establishing mitochondrial network topology and may be mechanistically distinct from DLP-mediated fission. Fenestrated sheets have also been observed in bloodstream form (BSF) *T. brucei* and are thought to resolve into tubules [58]. In addition, branching of the mitochondrion increases just prior to cell division, which may facilitate symmetrical division of the network [50, 58]. Whether these events are mediated by proteins that are conserved in the eukaryotic lineage, or whether they involve novel proteins specific to kinetoplastids, remains to be discovered. Several proteins were initially reported to play a role in mitochondrial shape but were later shown to be components of the mitochondrial import machinery [59–62]. Future experiments will reveal if these proteins moonlight as determinants of mitochondrial structure or if these complexes are responsible for importing other essential shape components. In either case, dynamic branching activities may need to be held in balance with periodic membrane fusion events in order to preserve the proper topology of the mitochondrial network.

Fusion/fission dynamics seem unlikely to operate in mitochondrial quality control in an organism with a single organelle and a single mitochondrial nucleoid, since any portion of the mitochondrion that became separated from the network would likely not contain any mitochondrial DNA. These fragments would have to either re-fuse to the network or be destroyed by mitophagy. Although physiological mitochondrial fragmentation has not yet been observed, investigators have noted that overexpression of the human pro-apoptotic protein Bax in *T. brucei* caused fragmentation of the mitochondrial network that was dependent on TbDLP [30, 50, 63, 64]. This effect was reversible, indicating that mitochondrial fragments are capable of re-fusing to create a continuous network. In further support of the existence of mitochondrial fusion in kinetoplastids, a mitofusin-like protein, TbMFNL, has been described [65]. Knockdown of this protein results in a highly fenestrated network in *T. brucei*, perhaps indicating that excess branches would typically be resolved through fusion. Furthermore, the mitochondrial duplication cycle for BSF *T. brucei* involves fusion of parts of the network [50, 58]. The presence of fusion indicates that there must be an opposing process of fission (or an alternative, branching activity), one that is not necessarily restricted to final division of the network. If both fusion and fission occur, they must be balanced in order to maintain mitochondrial shape, which would indicate that mitochondrial dynamics in kinetoplastids is not so different from other eukaryotes as previously thought.

It is possible that mitochondrial dynamics in kinetoplastids serves to homogenize mitochondrial components by mixing membranes and fluid compartments from different parts of the cell. This process may also allow for faster transport of mitochondrial components from one part of the organelle to another, a function that could be critical in organisms with a single large mitochondrion. In addition, the localized arrangement of mitochondrial DNA in kinetoplastids may present a unique challenge for the distribution of mitochondrial gene products within the organelle. In other eukaryotes, mitochondrial genomes are circular and found throughout each mitochondrion. In kinetoplastids, all copies of the mitochondrial genome are topologically interlocked and found in a single structure (kDNA) [66]. The kDNA is physically attached to the basal body via a transmembrane system of filaments called the tripartite attachment complex [54]. As the flagellar basal bodies serve to organize cell division, essentially acting as the cell’s centrosomes, their attachment to the replicated kDNA ensures the faithful distribution of this complex structure into daughter cells. The tripartite attachment complex will also physically restrict movement of the kDNA, retaining it in the kDNA pocket region. We know from epitope-tagging studies that mitochondrial ribosomes are found throughout the organelle [67], meaning that mitochondrial transcripts may need to be transported from the region of the kDNA to the rest of the mitochondrion for translation. Fusion of distant parts of the network may accelerate this transport beyond the rates of diffusion. Mitochondrial dynamics may therefore be conserved in kinetoplastids in order to fulfill specific requirements related to 1) having a single mitochondrion and 2) having all mitochondrial genomes collected at one end of the organelle. If this is the case, this process would be found in all kinetoplastid organisms with metabolically active mitochondria and would be required throughout the cell cycle.

As parasites, most kinetoplastids need to rapidly adapt their metabolism to changing nutrient availabilities. Mitochondrial remodeling is a well-studied part of metabolic adaptation during the life cycle of *T. brucei*, during which mitochondrial shape and activity change dramatically. Long slender BSF cells are highly glycolytic and have an unbranched mitochondrion, while developmental stages in the tse-tse fly, including procyclic form, rely on mitochondrial energy production and have a highly-branched mitochondrion [55, 56, 68]. Even within different compartments of the same host, the mitochondrion of *T. brucei* may need to adapt to different metabolic requirements. A recent study provides evidence that the metabolically downregulated BSF mitochondrion has a surprisingly complex proteome and could therefore be readily activated [69]. While this feature may facilitate differentiation to forms pre-adapted to life in the fly, the authors also point to recent discoveries of viable *T. brucei* cells in the skin and adipose tissues [70, 71]. Survival in these compartments may also require metabolic adaptation and alterations in mitochondrial shape.

*C. fasciculata* is most likely taken up by mosquitoes from a variety of habitats which could include pools of water and the surface of fruits or other plants. To survive, *C. fasciculata* parasites may also have to adapt their metabolism to make the most of their available nutrients. While we did not observe dramatic differences in mitochondrial structure or dynamics between nectomonads, which are more prevalent in the environment, and haptomonads, which are the primary form found in the mosquito [72], it should be noted that we generated these forms in culture in nearly identical nutrient environments. Whether mitochondrial shape and metabolism in *C. fasciculata* change in response to environmental conditions is an important question that could shed light on the conservation and possible functions of mitochondrial dynamics in a variety of parasites.

By imaging live cells, we were able to observe mitochondrial division and, indirectly, kDNA division. Division of the mitochondrion in *C. fasciculata* was closely coordinated with cytokinesis, as has been observed for *T. brucei* [30, 50, 58]. However, the mechanism of kDNA division is different between these two species. In all kinetoplastids, minicircles must be released from the catenated network prior to replication. They replicate as free circles, then are reattached at the antipodal sites flanking the disk. In *T. brucei* there is no relative movement between the kDNA disk and the sites of minicircle attachment. Minicircle removal from the center of the network and reattachment at the poles produces a dumbbell-shaped network which is essentially divided when the final minicircle is removed from the center for replication [73]. In contrast, kDNA in *C. fasciculata, T. cruzi, Leishmania*, and other species remain disk-shaped due to relative rotation between the disk and the reattachment sites, resulting in an even distribution of replicated minicircles [74]. In *C. fasciculata*, the kDNA divides in a single step mediated by an unknown enzymatic activity. By visualizing the kinetics of the loop of mitochondrion that contains the kDNA, we were able to image cells at the moment in which one kDNA network became two, shortly after the appearance of the cleavage furrow. Division occurred rapidly (under 10 minutes), making it difficult to imagine that the precise cutting and resealing of thousands of DNA circles occurred during this time. Possibly the two daughter kDNAs have already been divided at an earlier stage but remain tethered until the final division event. In some of our movies, an apparent release of tension occurs at the moment of division, propelling each daughter kDNA into its respective daughter cell with opposite trajectories. So far, we have been unable to find a DNA dye that fluorescently labels the kDNA in live *C. fasciculata* cells. Such a dye, or alternative methods to label the kDNA in live cells, would facilitate future studies of the process of kDNA division in *C. fasciculata*. Integration of the precise timing of kDNA division into our understanding of the mitochondrial duplication and division cycle provides a more complete picture of the highly-orchestrated cell cycle by which these organisms maintain their single-copy organelles.

Here we describe rapid mitochondrial dynamics in a model organism in which each cell contains a single mitochondrial network. Under certain conditions, yeast and mammalian mitochondria form a similar interconnected network [75], meaning that this mitochondrial configuration is not unique to kinetoplastids. Further studies on other kinetoplastid parasites will be needed to confirm that dynamics are a general phenomenon; however, we think it unlikely to be restricted to *C. fasciculata* since most features of the mitochondrial network are very similar between these related species. If mitochondrial dynamics is broadly conserved throughout eukaryotic evolution, study of this process in early-diverging eukaryotes such as kinetoplastids could provide important insights into its evolution and essential functions, while possibly explaining the metabolic adaptability of these parasites to different environmental conditions. Further, it is possible that intra-organelle fusion and fission reactions in *C. fasciculata* are mediated by novel or highly-diverged proteins, the study of which could reveal important mechanisms that will be applicable to the study of other eukaryotic model systems.

## Acknowledgments

This material is based upon work supported by the National Science Foundation under Grant No. MCB-1651517. The authors thank Stephen Beverley for providing the CfC1 parental cell line, Emmanuel Tetaud for providing the pNUS series of vectors and Stephen Hajduk for providing the mitoGFP construct. We also thank the PennVet Imaging Core facility for their expertise and the use of imaging equipment and software. This facility is supported by an NIH Shared Instrument Grant S10 OD021633-01 to Bruce Freedman. Thanks to Michael Povelones for comments on the manuscript and for help preparing figures.

## Supporting information

**S1 Fig.**
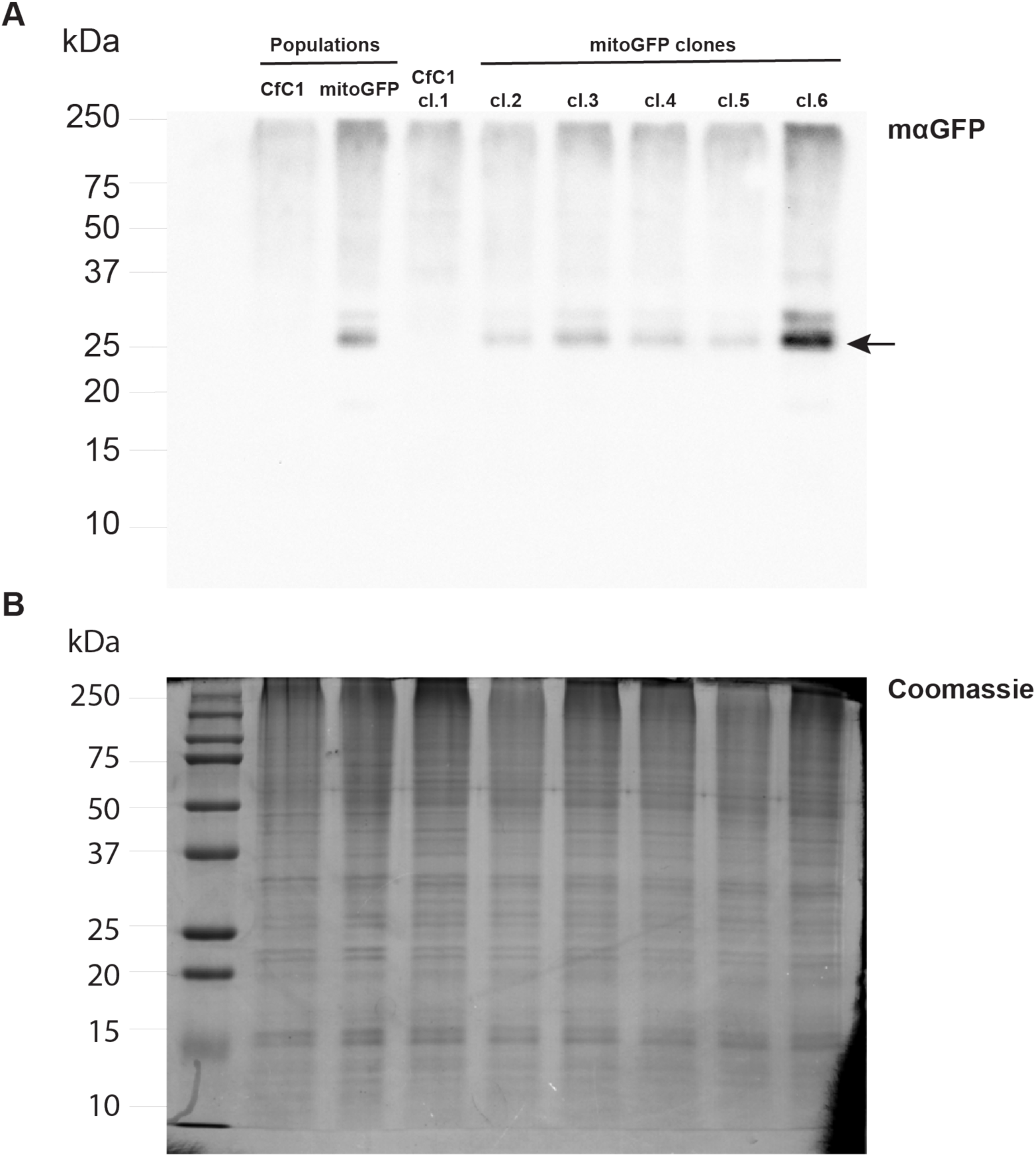
Clonal mitoGFP-expressing *C. fasciculata* lines were screened for GFP expression. A) Protein lysates from non-clonal (populations) and clonal lines were screened by western blot with anti-GFP antibody. Arrow indicates mitoGFP band. B) An identical gel as that used to generate the blot in A was run in parallel and stained with Coomassie as a loading control.

**S1 Movie. The mitochondrion is dynamic in *C. fasciculata*.** A representative swimming *C. fasciculata* cell expressing mitoGFP immobilized in agarose and imaged on a spinning disk confocal microscope. Maximum projections are shown including the frames shown in Fig 5A.

**S2 Movie. Mitochondrial dynamics in *C. fasciculata* nectomonads.** Swimming *C. fasciculata* cells expressing mitoGFP were immobilized in agarose and imaged on a spinning disk confocal microscope. Maximum projections of a representative cell are shown including the frames shown in Fig 5A.

**S3 Movie. Mitochondrial dynamics in *C. fasciculata* haptomonads.** Adherent *C. fasciculata* cells expressing mitoGFP were imaged on a laser scanning confocal microscope. Maximum projections of a representative rosette are shown and correspond to the frames shown in Fig 6A.

**S4 Movie. Dynamic fenestrated sheets appear in *C. fasciculata* mitochondria.** A representative adherent *C. fasciculata* haptomonad expressing mitoGFP was imaged on a laser scanning confocal microscope. Maximum projections are shown which have been color coded according to depth and which correspond to frames shown in Fig 6B.

**S5 Movie. Coordination of mitochondrial division and cytokinesis in nectomonads.** Maximum projection (deconvolved) of a swimming nectomonad *C. fasciculata* cell undergoing cytokinesis. The mitochondrion was imaged using mitoGFP. Time-lapse corresponds to frames shown in Fig 7A.

**S6 Movie. Coordination of mitochondrial division and cytokinesis in haptomonads.** A rosette of adherent *C. fasciculata* cells expressing mitoGFP. The cell at the bottom left is undergoing cytokinesis. Cleavage furrow ingression begins at 01:20 (mm:ss). Time-lapse of maximum projections corresponds to frames shown in Fig 7B.

**S7 Movie. Mitochondrial dynamics during cell division of haptomonads.** Maximum projection of a rosette of adherent *C. fasciculata* cells expressing mitoGFP. The cell at the right is undergoing mitochondrial division/cytokinesis. The top and bottom slices of the deconvolved Z-stack were removed in order to clearly visualize the division events.

**S8 Movie. Live-cell imaging of kDNA division in *C. fasciculata*.** Time-lapse of an adherent haptomonad *C. fasciculata* cell expressing mitoGFP. Several frames of the Z-stack have been removed from the maximum projections in order to clearly show the process of kDNA divison. Time-lapse corresponds to frames shown in Fig 7C.

**S9 Movie. The timing of kDNA division in *C. fasciculata*.** Deconvolved maximum projection of a *C. fasciculata* rosette expressing mitoGFP. The upper middle cell is in the initial stages of cytokinesis. The cell is oriented such that the anterior of the cell (where cleavage furrow ingression begins) is facing down. Division of the kDNA can also be observed.

